# Novel municipal sewage-associated bacterial genomes and their potential in source tracking

**DOI:** 10.1101/2021.07.15.452399

**Authors:** Blake G. Lindner, Brittany Suttner, Roth E. Conrad, Luis M. Rodriguez-R, Janet K. Hatt, Kevin J. Zhu, Joe Brown, Konstantinos T. Konstantinidis

**Author notes:** To whom correspondence should be addressed. Konstantinos T. Konstantinidis, 311 Ferst Drive, ES&T Building, Room 3321, Georgia Institute of Technology. Atlanta, GA, 30332. Department of Environmental Sciences and Engineering, Gillings School of Global Public Health, University of North Carolina at Chapel Hill, North, Carolina, NC 27599, United States.

## Abstract

Little is known about the genomic diversity of raw municipal wastewater (sewage) microbial communities, including to what extent sewage-specific populations exist and how they can be used to improve source attribution and partitioning in sewage-contaminated waters. Herein, we used the influent of three wastewater treatment plants in Atlanta, Georgia (USA) as inoculum in multiple controlled laboratory mesocosms to simulate sewage contamination events and followed these perturbed freshwater microbial communities with metagenomics over a 7-day observational period. We describe 15 abundant non-redundant bacterial metagenome-assembled genomes (MAGs) ubiquitous within all sewage inoculum yet absent from the unperturbed freshwater control at our analytical limit of detection. Tracking the dynamics of populations represented by these MAGs revealed varied decay kinetics, depending on (inferred) phenotypes, e.g., anaerobes decayed faster under the well-aerated incubation conditions. Notably, a portion of these populations show decay patterns similar to common markers, *Enterococcus* and HF183. Comparisons against MAGs from different sources such as human and animal feces, revealed low cross-reactivity, indicating how genomic collections could be used to sensitively identify sewage contamination and partition signal among multiple sources. Overall, our results indicate the usefulness of metagenomic approaches for assessing sewage contamination in waterbodies and provides needed methodologies for doing so.

## Introduction

Wastewater collection systems (or simply, collection systems) represent an important engineering control for the collection of human feces, commercial or industrial wastewaters, and sometimes stormwater, particularly in certain urban settings. The operation and maintenance of collection systems pose unique challenges, often due to their size, complexity, and capital costs (1–3). Population growth and distribution changes – especially growing urbanization trends – highlight the importance of maintaining and expanding efficient collection systems for an increasing fraction of the global population (4). Severe weather, pipe blockages, aging, and other issues of system failure can lead to the accidental release of untreated wastewater (sewage) from collection systems into waterways or floodwaters (1-3,5). As sewage is a significant reservoir of both chemical and biological pollutants, its release into the environment poses serious environmental and human health risks, including potential exposure to human pathogens (6–9) and possible dissemination of antimicrobial resistance genes (ARGs) among microbial populations (10–12).

Microbial source tracking (MST) refers to a collection of forensic tools developed to identify the presence and source of contamination among multiple probable fecal sources, including sewage (13). In large part, the technical approaches behind MST methods have been developed in response to both the difficulty of assaying for the diverse array of relevant human pathogens as well as the practical need to keep MST methods relatively rapid and inexpensive. Existing approaches have relied on indicator organisms to imply the presence of fecal pollution and sometimes as proxies for the presence of human pathogens in fecal contaminated waters. Specifically, fecal indicator bacteria (FIB) include an aggregation of bacterial populations considered representatives of microbial communities inhabiting the guts of warm-blooded animals. Widely used indicator organisms include *Escherichia coli* and *Enterococcus* spp. More recently, MST genetic markers from distinct bacterial lineages have been used that leverage known host specificity of distinct populations for source attribution (14). Some markers (e.g., HF183 assay targeting a human-associated *Bacteroides* clade) have found effective use in environmental management strategies as the basis for inferring the amount of sewage present and thereby, a potential array of pathogen concentrations for iterative risk assessment simulations (15). Yet, the use of FIB and MST gene markers has had challenges: most notably, that the concentration of most markers are rarely found to co-vary with pathogen concentrations, marker concentrations fluctuate with sewage age and the capability of FIB to adapt to environmental conditions can all combine to confound results interpretation (13,16–18).

Within recent years, targeted metabarcoding methods have examined sewage and sewage-contaminated waters via the 16S rRNA gene or the internal transcribed spacer (ITS) for prokaryotes and fungi, respectively (17,19–21). These studies have revealed a distinct sewage “microbiome” dominated by taxa that proliferate in collection systems, sometimes far beyond the abundance of human gut associated populations (22–24). However, these single-gene assays offer limited resolution to distinguish between environmental or non-environmental strains of the same species due to the high sequence conservation of the rRNA gene or the ITS region. Likewise, these methods do not provide information about the gene content associated with important populations (e.g., emergent pathogens, ARGs) or resolve finer community-wide compositional shifts (17, 25). Therefore, rRNA gene-based approaches are limited with respect to quantifying health risks associated with the detection of biomarkers or guide the development of more holistic environmental management criteria (e.g., site specific criteria).

Whole genome shotgun sequencing (or metagenomics), which recovers fragments of the genomes in a sample, have revealed that bacteria and archaea predominantly form sequence-discrete populations with intra-population genomic sequence relatedness typically ranging from ∼95% to ∼100% average nucleotide identity (or ANI) depending on the population considered (26). Metagenomic approaches offer unique advantages for environmental health monitoring tasks including: 1) extensive gene content information of abundant populations, 2) precise ecological estimates of relative abundance at the species level and 3) examination of intra-species diversity (27). Despite its potential for circumventing some of the challenges facing existing MST and metabarcoding methods, whole genome shotgun sequencing has not been fully utilized in monitoring municipal sewage pollution. To date, most applications have focused on understanding the microbiology of biological wastewater treatment, treated effluents and their receiving waters, or viral populations (12,28,29). In part, this is because it remains unclear how to best merge the methods and bioinformatics behind metagenomic practices with existing MST and environmental monitoring paradigms (30). Widespread application of this technology in the field requires that several outstanding issues be resolved, including the detection limits of metagenomic analyses, whether whole and/or metagenome-assembled genomes (MAGs) can serve as source-specific fecal contamination markers and how metagenomic approaches can infer the relative contribution of various fecal inputs (referred to hereafter as “source partitioning”).

Here, we offer a genome-centric view of sewage-related bacterial populations and explore their relationships with culture and PCR-based markers during a simulated failure of a collection system (e.g., an overflow event). Specifically, we simulated sewage contamination events in lake water obtained from a local drinking and recreational use reservoir, Lake Lanier (GA, USA), within dialysis bag mesocosms that were incubated in darkness for one week. Shotgun metagenomic sequencing was performed to search for potential sewage-specific biomarkers, test the effectiveness of genome collections for fecal source attribution and partitioning, and directly screen for both pathogens and antimicrobial resistance genes. Lastly, we propose a theoretical analytical limit of detection for metagenomics that could help guide the future application and interpretation of whole genome shotgun sequencing to these issues.

## Methods

### Sample collection, mesocosm setup, and sample processing

Samples were collected in sterile glass 1 L bottles from the primary influent of three WWTPs located in the Atlanta Metropolitan region of Georgia (USA) to serve as representatives of sewage across three different sewersheds. Each sewershed was comprised of collection systems with separate stormwater and wastewater conveyance (i.e., separate sewers). Approximately 50 L of surface water from Lake Lanier, Georgia was also collected concurrently. Hereafter, these sample groups are referred to as sewersheds A, B, and C. All sewage and water samples were immediately transported to the lab and stored in darkness at 4 °C until mesocosm setup, which occurred within 24 hours. For mesocosm setup, 40 L tanks were filled with lake water and a pump installed for aeration. Experimental dialysis bags were prepared with 110 mL 10% (v/v) sewage and lake water mixture and control bags were filled with 110 mL uninoculated lake water and closed on both ends using polypropylene Spectra/Por clamps (Spectrum Laboratories). Both experimental (n=12 x 3 sewersheds = 36 bags) and control (n=12 bags) dialysis bags were then added to the tank. A small headspace of air was left in each bag when sealing with clamps so that they could float freely in the tank. Dialysis bag pore sizes (6-8 kDa molecular weight cutoff) permit the transport of small molecules and ions, but bacterial and viral particles are contained within the bags. Mesocosms were kept in darkness at 22°C throughout the duration of the experiment. Sampling occurred at 1, 4, and 7 days by retrieving experimental and control bags from the mesocosm for destructive processing.

Mesocosm sampling, DNA extraction and subsequent qPCR analysis occurred as described previously in Suttner et al (35). Briefly, water samples were passed through 0.45 μm poly-carbonate (PC) membranes and stored at -80 °C in 2 mL screw cap bead tubes until processed (within 1-3 months). EPA Method 1600 (31) was followed for enumerating volumetric *Enterococcus* CFUs. DNA was extracted from PC membranes using the Qiagen PowerFecal kit and following the manufacturer’s instructions with only one exception: mechanical cell lysis was performed by bead beating in two 1-minute intervals using the Biospec Mini-Beadbeater-24 with icing between intervals. These DNA extractions were used for qPCR with the HF183/BFDRev assay (32) and a universal 16S rRNA gene qPCR assay (GenBac16S) to quantify 16S rRNA gene copies across samples (34). Metagenomic sequencing was performed using the Illumina Nextera XT kit with library average insert size determined on an Agilent 2100 instrument using a HS DNA kit and library concentrations determined using the Qubit 1 X dsDNA assay. Samples were then pooled and sequenced on the HiSeq 2500 instrument as described previously (33).

All qPCR reactions were run using an Applied Biosystems 7500Fast thermocycler and the cycling parameters were as follows: 2 min at 50 °C, 10 min at 95 °C, and 40 cycles of 15 sec at 95 °C and 60 sec at 60 °C. Assay reactions used 2 μL of template DNA in 20 μL qPCR reactions with the TaqMan Universal PCR Master Mix (Applied Biosystems). The primer and probe concentrations were 0.25 µM for HF183 assay and 0.3 µM for the GenBac16S assay. Template DNAs were run undiluted or diluted 5-fold (to remove the effect of PCR inhibitors) depending on the expected marker concentration and quality of each sample. Further details on qPCR reaction set up and standard plasmids for absolute quantification are provided in Suttner et al. (35) and reiterated within the supplement here (**Supplement Table S1**). To test for extraneous DNA and potential contamination from sample handling, 50 mL of sterile PBS was also filtered onto PC membranes and processed following the same DNA extraction at every sampling time point as described above.

**Table 1.**
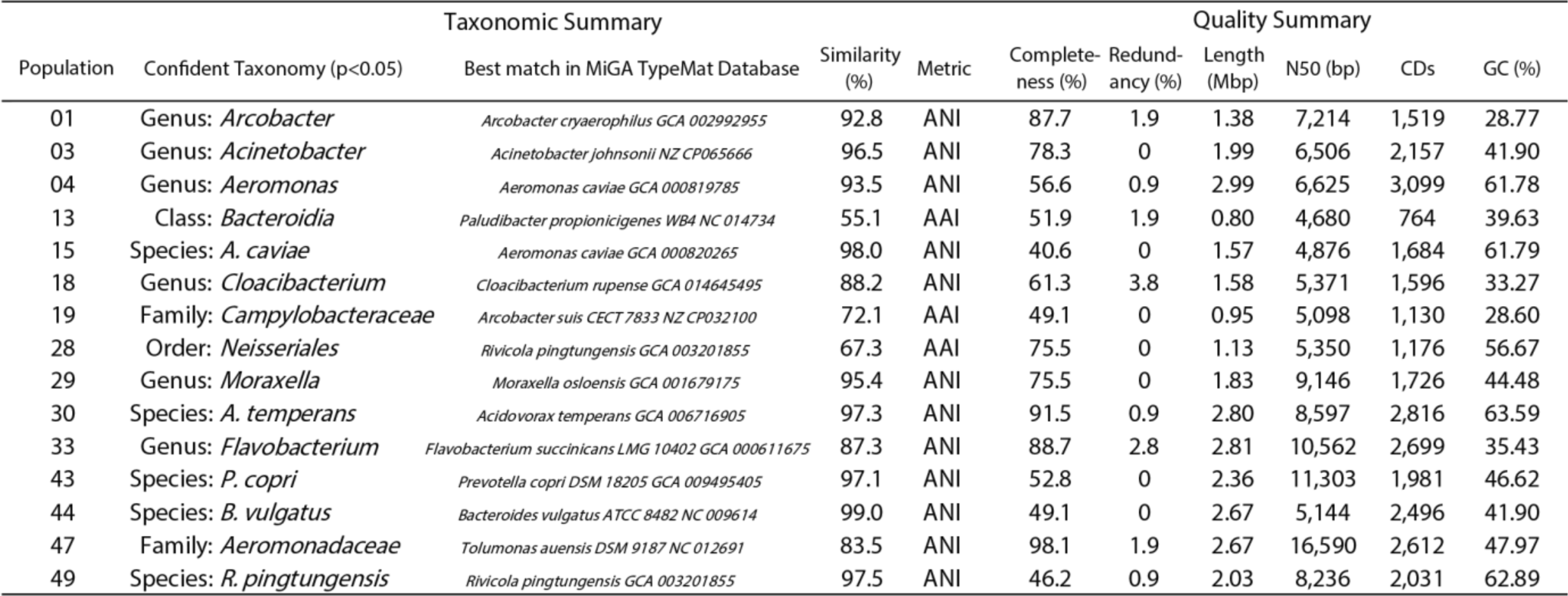
Summary of representative MAGs recovered in this study representing sewage-associated populations.

### Bioinformatics sequence processing and population genome binning

Short reads were quality trimmed and Nextera adapters removed with Trimmomatic 0.39 (36). Quality trimming was performed to remove poor quality bases along both ends of sequences and subsequent removal of any sequences below 50 bp in length. *k*-mer based operation of Nonpareil 3.304 (-T kmer) was used to estimate the fraction of alpha diversity covered by the sequencing effort of each metagenome (37). Beta diversity across trimmed short reads was assessed with the default settings of simka 1.5.1 based on Bray-Curtis dissimilarity values and visualized by principal coordinate analysis (PCoA) (38). Kraken2 was used to assign taxonomy and estimate simple relative abundance against a custom library, including bacteria, archaea, viruses, protozoa, human, and fungal reference genomes at the rank of class (39). Trimmed short reads were assembled individually with IDBA (UD) 1.1.3 and SPAdes (“--meta”) 3.14.0 using *k*-mer sizes between 20 and 127 (40, 41). Contigs shorter than 3Kbp were removed prior to population genome binning, which was performed with MaxBin 2.2.7 and MetaBAT 2.12.1 (42, 43). Additionally, in a parallel workflow, trimmed short reads were normalized via the BBNorm function of the BBtools suite (version 38) to bring read depths between 10-30X sequencing depth and then subsequently assembled and binned as described above (44). All resulting metagenome-assembled genomes (MAGs) from both regular and depth-normalized short read assemblies were dereplicated using MiGA 0.7.24.0 via the derep_wf function (45). Groups of MAGs sharing ANI ≥ 95% were clustered into species-like populations (hereafter, “populations”) with representative MAGs for each population selected by highest completeness and lowest redundancy. Populations with no representative MAG having a MiGA quality score above 30% and/or redundancies below 5% were excluded from further analysis. Both Traitar 1.1.2 and MicrobeAnnotator were used with default settings to infer potential phenotypes and annotate draft genomes, respectively (46, 47). Lastly, MAGs were screened for cross-reactivity using the FastANI tool to search for other genomes with ANI ≥ 95% across a suite of reference databases (48).

### Annotation of sequence data

From the PATRIC database, version 3.6.9, 1097 pathogenic bacterial genome accession IDs were recovered by querying for host name “Human, homo sapiens” and “good” quality. This included both genomes tagged as “Reference” (n=28) and “Representative” (n=1069) (49). Of these, 1076 genomes were recoverable from NCBI for use in this study. Abundance estimates of pathogen genomes were assessed by short read mapping with Magic-BLAST 1.4.0 (-splice F) (50). Resulting alignments were filtered using minimum cut-off of 70 bp alignment length, 95% query coverage by alignment and 95% identity to avoid spurious matches. Additionally, for virulence gene detection, only experimentally verified nucleotide entries in the Virulence Factor Database (51) were used. Evaluating MAG abundance across the time series was accomplished similarly using Magic-BLAST 1.4.0, where MAGs were concatenated into a single library to which reads were competitively mapped. Additionally, DIAMOND 2.0.1 (blastx --ultra-sensitive) was used to search short reads against the reference gene sequences of pre-compiled 150 bp β-lactamase ROCker models to reliably identify short reads belonging to β-lactamase encoding genes (52, 53). Reads mapping to these reference sequences were selected for best bit-score alignment and subsequently filtered by ROCker v1.5.2 as described previously (54).

### Estimation of limit of detection, and relative or absolute abundance

For a reference genome, MAG, or gene to be considered detected in a sample, at least 10% of the target sequence was required to be covered by reads (i.e., breadth of coverage: hereafter, C), as proposed previously for robust detection of targets in metagenomic datasets (55). Or, as written, the analytical limit of detection (LOD) used here:

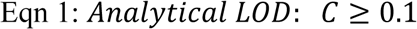

The LOD was automatically implemented by calculating sequencing depth and breadth similarly to Rodriguez-R et al (56) for estimating “Truncated Average Depth” at 80% (hereafter, the function TAD80). Python scripts used for this approach are available online at: https://github.com/rotheconrad/00_in-situ_GeneCoverage. In short, the TAD80 function estimates sequencing depth by first sorting genomic positions according to their sequencing depth and then removing the upper 10% and lower 10% of positions before averaging the sequencing depth along the remaining 80% of positions. Since truncation of targets with breadth of coverage near the detection limits (e.g., C ≈ 0.1) could introduce artificially lower values, a quantification threshold was also necessary to avoid systemic underestimation of abundance for targets near LOD. From Lander and Waterman (57), breadth of coverage (C) is related to sequencing depth (ρ) by the following:

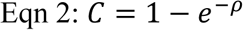

Thus, for the analytical LOD defined above, the expected sequencing depth (ρ) is simply -ln(0.9) for targets at detectable limits. We formalize a quantification threshold which measures whether a target is quantifiable following application of the truncation function (TAD80) with:

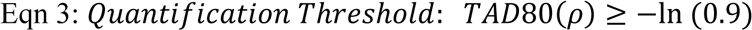

For simplicity in our metagenomic results, we describe those targets which satisfied the LOD condition but were below the quantification threshold as targets that were “detected but not quantifiable” (DNQ).

To convert coverage of detected target genomes to absolute abundances (e.g., cells/mL), the following approach was used. Single copy gene coverage or genome equivalents (GEQ) and average genome size (AGS) of metagenomes were evaluated using MicrobeCensus 1.1.0 (58). The 16S rRNA gene-carrying reads were identified and extracted using sortmeRNA 4.2.0 and the average 16S rRNA gene coverage was estimated as the sum of extracted read lengths divided by 1540 bp, the average length of the bacterial 16S rRNA gene (59, 60). Average 16S rRNA gene copy number (16S ACN) for each metagenome was determined by the ratio between 16S rRNA sequencing depth (ρ*_16S_*) and GEQ:

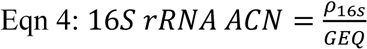

The copy number of the 16S rRNA gene per mL as quantified by qPCR was divided by the 16S rRNA ACN to obtain an estimate for the number of cells in each sample, assuming that one prokaryotic genome was approximately equivalent to one prokaryotic cell:

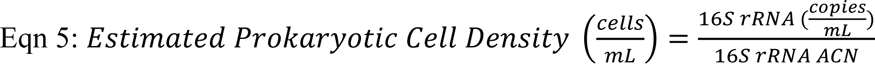

These measures were taken to help control for bias in relative abundance estimation due to changes in overall microbial load (cells per volume) and 16S rRNA gene ACN variation throughout the experiment (61, 62). Finally, absolute abundances were estimated by multiplying a population’s genome equivalents by the estimate for the number of cells in a sample. This was accomplished using the following equation for a given population via the truncated average sequencing depth [TAD80(ρ*_i_*)], GEQ and total estimated prokaryotic cell density:

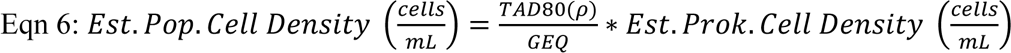

Further, an extension of our definitions of LOD was used in tandem with cell density estimations for theorizing the smallest abundance detectable as a function of GEQ and cell density via:

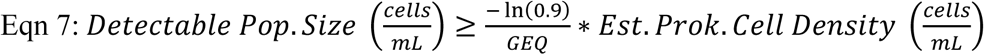

## Results

### Culture and qPCR Data

Both fecal indicators (*Enterococcus* and HF183) were in the same order of magnitude across the sewage samples gathered as inoculum for the mesocosms. Sewage from sewersheds A and B contained counts with averages of 3.7E+04 and 3.1E+04 Enterococci CFUs/100mL and 2.4E+06 and 3.6E+06 HF183 copies/mL, respectively. Within sewershed C, counts were lower having 1.3E+04 Enterococci CFUs/100mL and 1.5E+06 HF183 copies/mL. Similarly, quantification of the 16S rRNA gene copy number within the inoculum indicated that overall, microbial loads were lower in sewershed C than sewersheds A and B (**Supplement Figure S1**). Monitoring Enterococci and HF183 qPCR markers across the mesocosm timeseries revealed that the markers decreased throughout the experiment in all replicates but were still detectable at day 7 and remained higher than the established or recommended water quality criteria for recreational use waters (i.e., 36 CFUs/100mL and 41 HF183 copies/mL). Only the HF183 marker within sewershed C mesocosm decreased below detection on Day 7 (**Figure 1**). Neither marker was detected in the (un-inoculated) freshwater sample serving as control at any time point during mesocosm operation.

**Figure 1.**
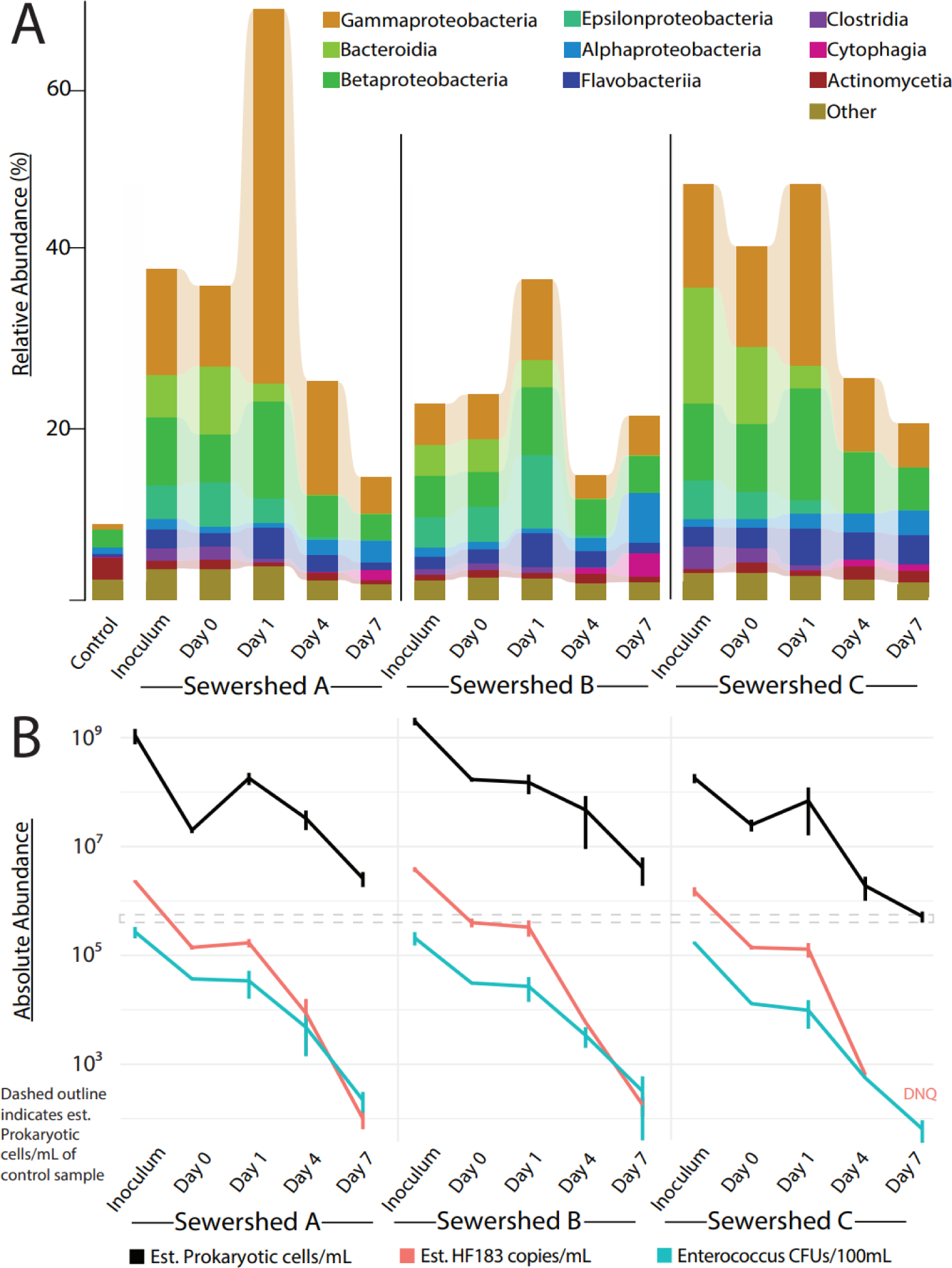
**Panel A**: Class level abundances across control, inoculum and timeseries for sewersheds A, B and C based on kmer classification by Kraken2 against a custom-built database of reference genomes. Total height of bars represents the percentage of kmers confidently classified to the corresponding taxon (Figure key). The maximum and minimum percentages of kmers confidently classified were 69.0% from Sewershed A day 1 and 8.9% from the control, respectively. **Panel B**: Estimated cell density, estimated HF183 copy concentration and Enterococci colony forming units (CFU) for the same samples. The dashed lines indicate the estimated cell density range for the control sample. HF183 was detected but not quantifiable (DNQ) for Sewershed C on day 7.

### Estimated Microbial Load

Prokaryotic cell density of the inoculum varied per mesocosm based on quantification of the 16S rRNA gene (see Methods for details): 1.1E+09, 2.0E+09, and 1.8E+08 cells/mL were estimated for sewersheds A, B and C, respectively. Following dilution and mixing of the inoculum into the mesocosms, day 0 estimates for cell densities were 2.0E+07, 1.7E+08, and 2.5E+07 cells/mL. Thereafter, cell density in both sewershed A and sewershed C mesocosm increased considerably in the first 24 hours to 1.8E+08 and 6.9E+07 estimated cells/mL (a 924% and 275% increase) while sewershed B decreased to (an estimated) 1.5E+08 cells/mL. Subsequent time points revealed steady decreases in cell densities approaching the control cell density at day 7 of 7.9E+05 estimated cells/mL (**Supplement Table S2**).

### Metagenomic-based Coverage and Compositional Shifts of the Mesocosms Over Time

Between 1.5 Gbp to 3.5 Gbp of data per sample remained following read quality trimming and adapter removal, which corresponded to a range of 9 to 27 million reads. Sequencing effort covered between 36 to 67% of expected nucleotide diversity (*N_d_*) across all samples based on the Nonpareil algorithm, which estimates sequence coverage based on the degree of redundancy among the metagenomic reads available for each dataset (36). This level of coverage is adequate for comparing the abundance of features (e.g., genomes, genes) across samples (63). *N_d_* estimations of the inoculum and control samples were similar, and day 0 values closely followed that of their respective sources. A decrease in *N_d_* occurred within the first 24 hours for all three biological replicates; lower diversities were observed in day 1 samples compared to those for the inoculum, day 0 samples and the control. The sewershed B series increased in diversity for the remaining days while both sewersheds A and C vacillate thereafter (**Supplement Table S2**).

Observations of beta diversity revealed that the earlier timeseries samples (day 0 and day 1) remained quite similar to the inoculum. By day 4, considerable shifts in community composition were observed driving the sewage contaminated waters closer to the control (**Supplement Figure S2**). *k*-mer mapping to characterize these community-wide shifts using Kraken2 at the class level showed the depletion of *Bacteroidia*, *Epsilonproteobacteria,* and *Clostridia* following inoculation. None of these classes were detectable in the control samples. An increase of *Gammaproteobacteria* abundance occurred within the first 24 hours across all replicates after which this class gradually decreased in abundance with time. Additionally, increases in *Alphaproteobacteria* and *Cytophagia* occurred in later time points (day 4 and day 7), far beyond the increase observed in the control, suggesting that the later timepoint samples had not yet fully recovered from perturbation. Class level relative metagenome-based abundances, qPCR, culture, and cell density estimation results are summarized on **Figure 1**.

### Sewage-associated Population Genome Binning

Seven hundred twenty MAGs were recovered from inoculum and timeseries sample assemblies. The 720 MAGs were dereplicated at the ANI ≥ 95% level, resulting in 49 MAGs representing sequence discrete populations (hereafter, simply “populations”). Competitive read mapping to the representative MAG of these populations revealed two groupings delineated by their presence or absence in the inoculum. Of the total 49, 33 populations were detected within sewage inoculum samples with varying degrees of prevalence across replicates. We selected a subset of 15 of these 33 populations that were above the quantification threshold in each inoculum sample, which we refer to as “sewage-associated populations”. This selection process was motivated twofold: First, to focus only on core populations shared between the inoculum recovered from each sewershed examined herein. Second, as an effort to exclude potentially noisy, nonspecific, or transient populations from further analysis. The sewage-associated populations and their representative MAGs are summarized in **Table 1**. Additionally, we validated our analytical detection and quantification limits using mock data of known composition to ensure these criteria were suitable for identifying sewage-associated populations (**Supplement Table S3.A**) (64). We found our approach, as described in Methods (Eqn. 1 and 2), was robust for reducing quantification error and detected targets of known relative abundance as expected according to sequencing effort and target genome size, except on very limited occasions when close relatives were present in the sample at frequencies many times greater than the target genome. (**Supplement Table S3.B,C**).

Our collection of ubiquitous sewage-associated populations in sewersheds A, B, and C inoculum metagenomes were represented by respectively 9.5%, 5.7%, and 13.3% reads and 15.9%, 8.8%, and 19.6% of GEQ. Estimated absolute abundances of these populations varied across the samples, from a maximum of 4.4E+07 cells/mL (Pop.01, sewershed B) to a minimum of 2.3E+05 cells/mL (Pop.04, sewershed C). Within the inoculum, the median and mean absolute abundances of an individual sewage-associated population was 5.3E+06 and 8.4E+06 cells/mL, respectively. Overall, sewershed C had substantially lower population densities due to the difference in total microbial load compared to sewersheds A and B, as noted above. Consistently, the sewage-associated populations presented here capture a larger portion of the metagenomic samples associated with sewershed C (compared to A or B), further indicating that the sewershed C samples may have simply had more dilute microbial load at the time of sampling. Overall, these results reveal that this collection of populations consistently represent highly abundant members of the sewage microbiome across biological replicates and a substantial part of the total sewage microbial community.

Comparison of the corresponding representative MAG sequences against type material in the MiGA “TypeMat” database (45) revealed several entries with close matches to previously described taxa at the species level (e.g., >95% ANI) including *Aeromonas caviae* (Pop.15), *Acidovorax temperans* (Pop.30), *Prevotella copri* (Pop.43), *Bacteroides vulgatus* (Pop.44), and *Rivicola pingtungensis* (Pop.49). Of the remaining, six populations matched known genus representatives, potentially representing a novel species of the matching genera. Two populations matched members of a known family, one to members of a known order, and one to members of a known class (**Table 1**). The population with the most distant match in the database (Pop.13, matching class *Bacteroidia*) with 55.1% average amino acid identity (AAI) to *<u>Paludibacter propioncigenes</u>*.

Collections of bacterial isolate genomes and/or MAGs from freshwater (56), activated sludge (65), anaerobic digestors (66), the human gut environments (67), and the broad general-purpose GEMs catalog (68), were examined to assess specificity between these 15 sewage-associated populations and other microbiomes. Of these sewage-associated populations, some (n=11) may belong to species with members also inhabiting non-sewage microbiomes such as biological wastewater treatment processes or the human gut (**Supplement Table S4**). Importantly, only a single population, *Moraxella* (Pop.29), was found via these database searches to match (95.1-95.0% ANI, borderline of universal species cutoff) genomes recovered from aquatic environments (both marine and freshwater) (47). This finding suggests Population 29 could be less effective as an entry in a sewage-specific genomic library utilized for MST approaches if other *Moraxella* are in high abundance within the pristine environment.

### Sewage-associated Population Decay and Putative Phenotyping

Overall, all populations experienced rapid decline in estimated cell densities across the timeseries with most populations below detection limits following day 4. *Acinetobacter sp.*, *Cloacibacterium sp*., *Acidovorax temperans*, and *Flavobacterium sp*. (Pop.03, Pop.18, Pop.30 and Pop.33, respectively) were detectable in at least one biological replicate at day 7 but most of these observations were below quantification. Signal from sewershed A had the greatest persistence; of the four mesocosms with quantifiable levels of a sewage-associated population by day 7, three belonged to the series of sewershed A. Notably, *Acidovorax temperans* (Pop.30) was the only population detected at day 7 in all three sewersheds (**Figure 2**).

**Figure 2.**
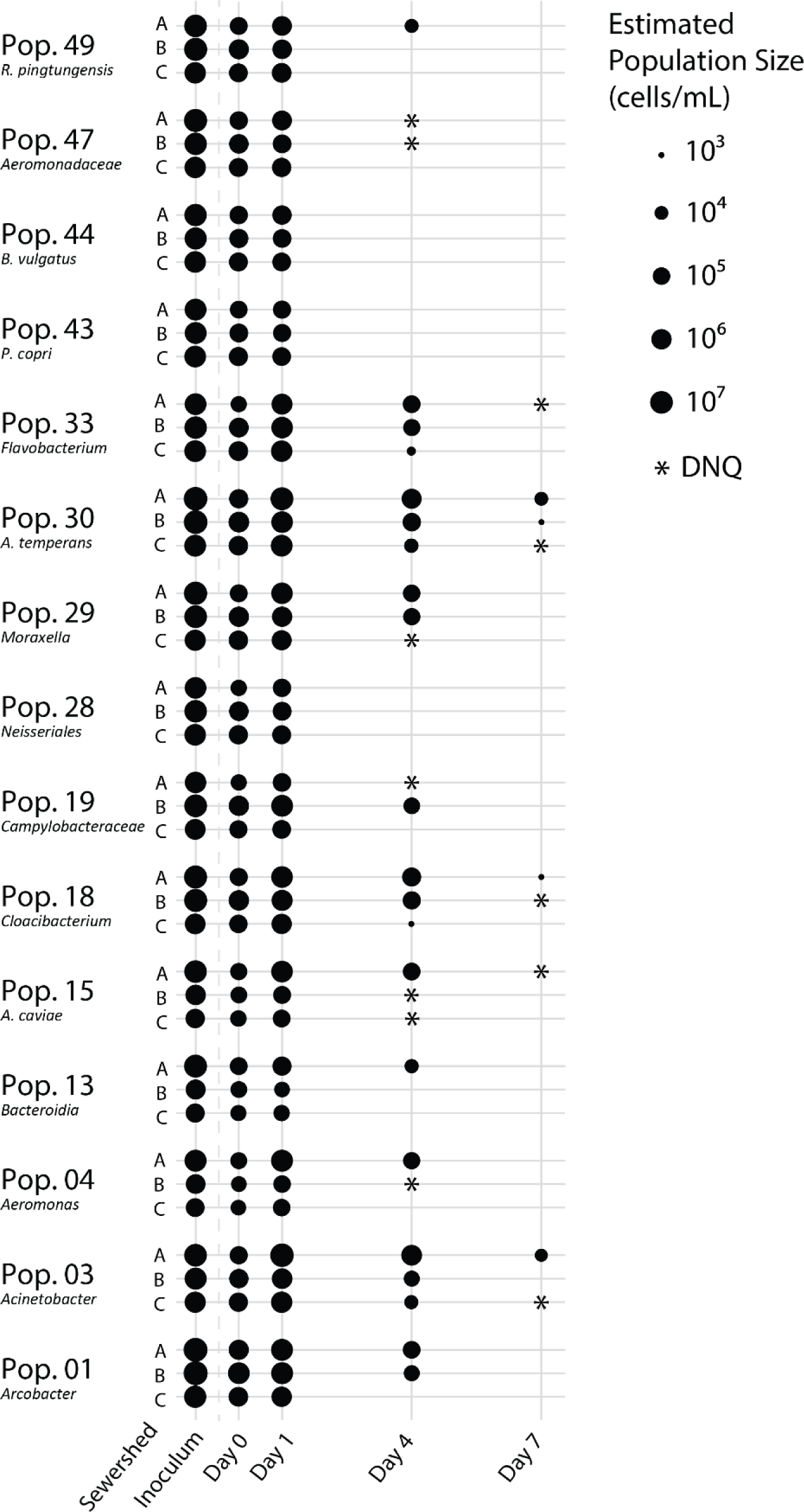
Estimated cell densities of sewage-associated populations across inoculum and timeseries samples. Cell densities (absolute abundances) were estimated as described in the Materials and Methods section.

All populations remaining detectable at day 7 were putatively phenotyped as aerobic or facultatively anaerobic by Traitar analysis except for *Cloacibacterium sp.* (Pop.18), which could not be confidently classified. Nonetheless, *Cloacibacterium sp.* belongs to a genus of facultative anaerobes (*Cloacibacterium*), suggesting that it likely is a facultative population and that the representative MAG did not contain the necessary genes for confident phenotyping due to incompleteness. No population – regardless of (predicted) preference for oxygen – showed an increased estimated cell density outside the first 24 hours of the incubation. All sewage-associated populations were likely gram negative, rod or oval-shaped bacteria as predicted by Traitar (**Supplement Figure S4**).

MicrobeAnnotator indicated *Acinetobacter sp.* (Pop.03) and *Acidovorax temperans* (Pop.30) contained modules for aromatic carbon degradation which were rare genomic features among the representative MAGs. *Acinetobacter sp.* (Pop.03) was reported to contain complete benzoate degradation and catechol ortho-cleavage modules, while *Acidovorax temperans* (Pop.30) contained complete catechol ortho-cleavage and incomplete catechol meta-cleavage modules but none for benzoate degradation (**Supplement Figure S5**).

### Human Markers vs Sewage-associated Populations

Our results suggested that several of the sewage-associated populations are possibly linked to the human gut microbiome (**Supplement Table S4**). Based on AAI values, Pop.43 and Pop.44 were assigned to *Bacteroidales* lineages but likely represent a different lineage than that represented by HF183 based on the 16S rRNA genes carried on these populations’ closest complete genome matches from a cultured representative (See **Table 1**). Modelling the linear relationship between either HF183 or *Enterococcus* concentrations against the estimated cell densities of the sewage-associated populations revealed divergent results for both markers. HF183 had excellent correlations against some populations (i.e., anaerobic Pop.43 and Pop.44, and aerobic Pop.30 and Pop.28) but highly variable correlations overall (R^2^ between 0.35 to 0.97) while *Enterococcus* had worse correlations but with a tighter range (R^2^ between 0.5 to 0.8) (**Figure 3**). As noted above, not all the sewage-associated populations highlighted as potentially co-habiting the human gut co-varied in abundance as well with HF183 concentrations. For example, correlations with HF183 concentrations were moderate with the presumed aerobes of Pop.03 (R^2^ = 0.69) and Pop.29 (R^2^ = 0.75) but poor for the facultative Pop.15 (R^2^ = 0.35).

**Figure 3.**
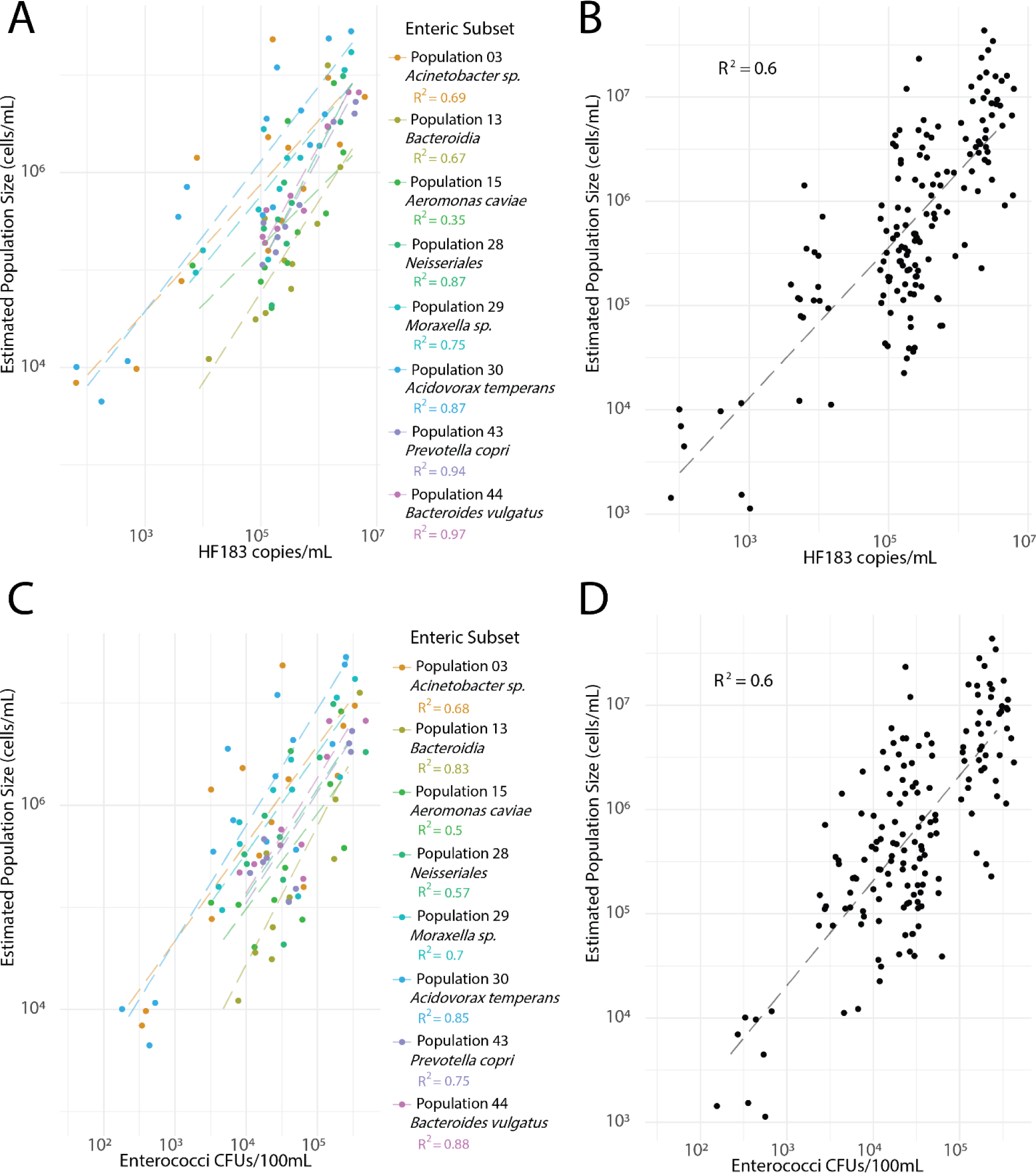
Log-log scatter plots of estimated population densities across inoculum and timeseries samples against HF183 and Enterococci concentrations. Lines of best fit are shown dashed with their associated coefficients. Panel A: HF183 copy number versus the concentration of sewage-associated populations likely to also be enteric (n=8). Panel B: HF183 copy number versus the concentration of all sewage-associated populations (n=15). Panel C: Enterococci concentration versus the concentration of sewage-associated populations likely to also be enteric (n=8). Panel D: Enterococci concentration versus the concentration of all sewage-associated populations (n=15).

### Genome-based Source Tracking

We aimed to demonstrate how read mapping metagenomic data to genome collections can perform differential source attribution of the fecal contamination within our single input mesocosm experiments. Our goal was to construct libraries of genomes representing the microbial pollutants expected to belong to a specific fecal source. To accomplish this, we downloaded MAGs, isolate genomes, and other reference genomes from several largescale studies of host microbiomes to create a library for determining source attribution with whole-genome sequences (67, 69–71) The collected data included genomic entries representing the fecal microbiomes of humans (n=4644), pigs (n=1667), and chickens (n=5675) and the rumen microbiome of cows (n=2124). Using MiGA, we dereplicated entries from each collection to obtain representative genomes with the highest quality for groups of genomic entries with ANI ≥ 95% to each other to reduce both the size of the dataset and redundant entries representing highly similar populations within a collection of host-associated genomes. This approach was the same as described in our methods for processing and selecting the MAGs described above and shown on **Table 1**. Using these dereplicated genomes and including the 15 representative MAGs we produced herein, we searched for and removed any instances of ANI ≥ 95% matches across these collections of host-associated genomes to control for potential cross-reactivity. One exception for which we did not remove matching genomes was between matching human and sewage genomes since they represent the same fecal source and would not complicate interpretation of results (**Supplement Table S4**). Our library construction and curation efforts are summarized in **Figure 4**.

**Figure 4.**
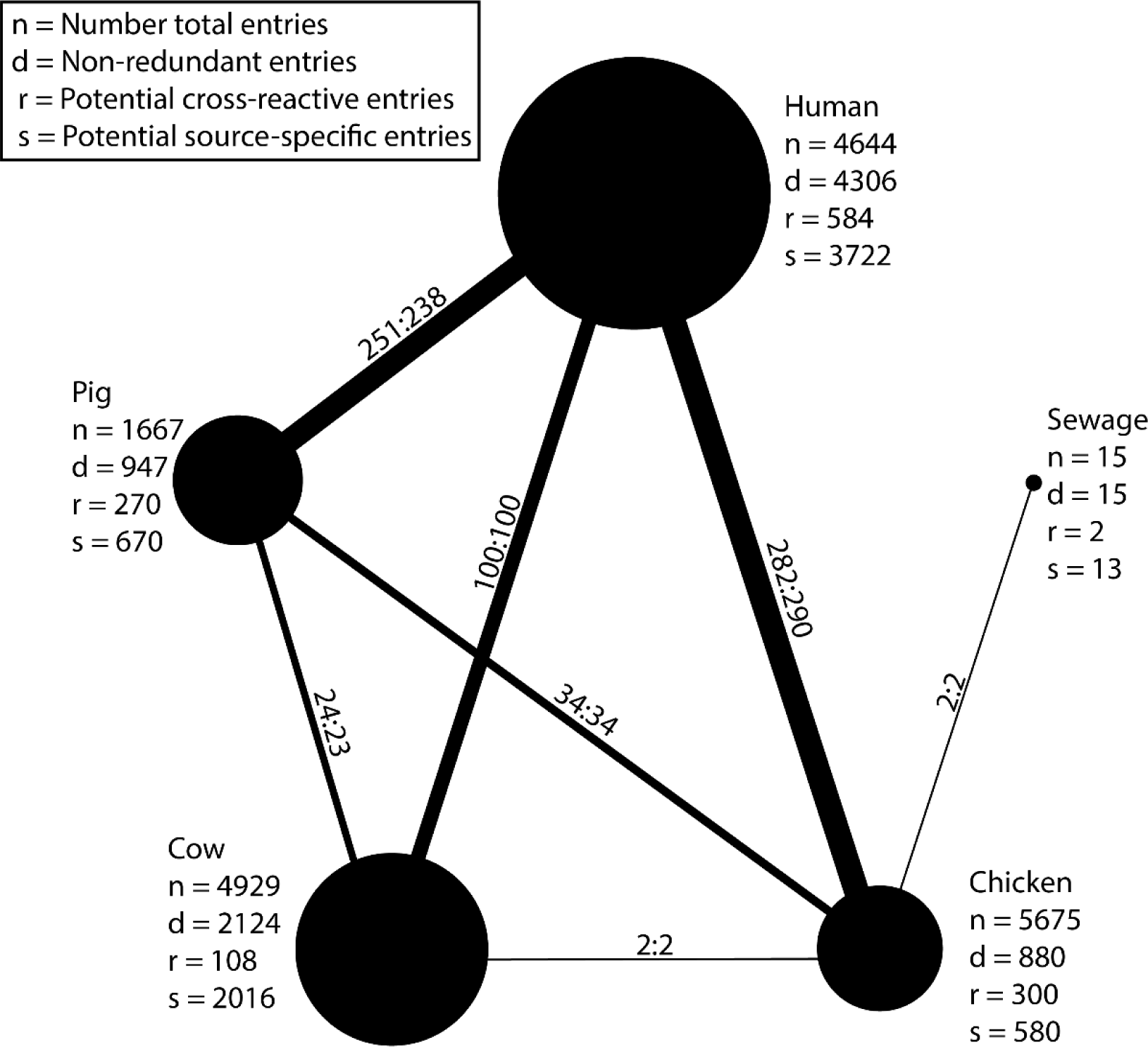
Overview of the curated genomic library for source attribution. Nodes represent sets of genomes recovered from public datasets of a given host microbiome, either “human”, “pig”, “cow”, and “chicken”. The “sewage” genomes shown here are those MAGs produced in this study. The radius of a node is proportional to the square root of the number of dereplicated genomes (d) remaining in the set following processing as described in the Methods section. Edges connecting nodes represent the amount of potentially cross-reactive genomes at the ANI ≥ 95% threshold level, and the line weights are drawn proportional to the square root of the largest number of these potentially cross-reactive matches (r). Ratios between nodes represent the number of matching genomes, with the value nearest a particular node representing the number of genomes from that dataset which matched across libraries.

Next, we aimed to use our host-specific source libraries to perform source attribution and partitioning as if our mesocosm data represented metagenomes recovered from a waterbody contaminated by a single unknown source. Thus, we performed competitive read mapping of the metagenomic data to the finalized non-redundant genomic library using Magic-BLAST and custom scripts for TAD80 calculation as described above for tracking the sewage-associated populations individually. Resulting TAD80 values were summed within each source category and normalized to GEQ to allow interpretation of these results as the percentage of contribution from each fecal source in the form of % GEQ (e.g., percentage of prokaryotic genomes) belonging to a source (**Figure 5, A**). No source category was detected in the control samples and signal from our collection of sewage MAGs dominated the timeseries across all sewersheds but rapidly disappeared after day 4. The combined human signal followed a similar pattern as the sewage though usually at about 10% less GEQ. The pig, cow, and chicken source categories were usually not detected or were consistently <0.1% GEQ. Hence, this approach provides the means to assess contamination at the metagenomic read level, circumventing a substantial computational burden to assemble, bin and obtain de-replicated MAGs.

**Figure 5.**
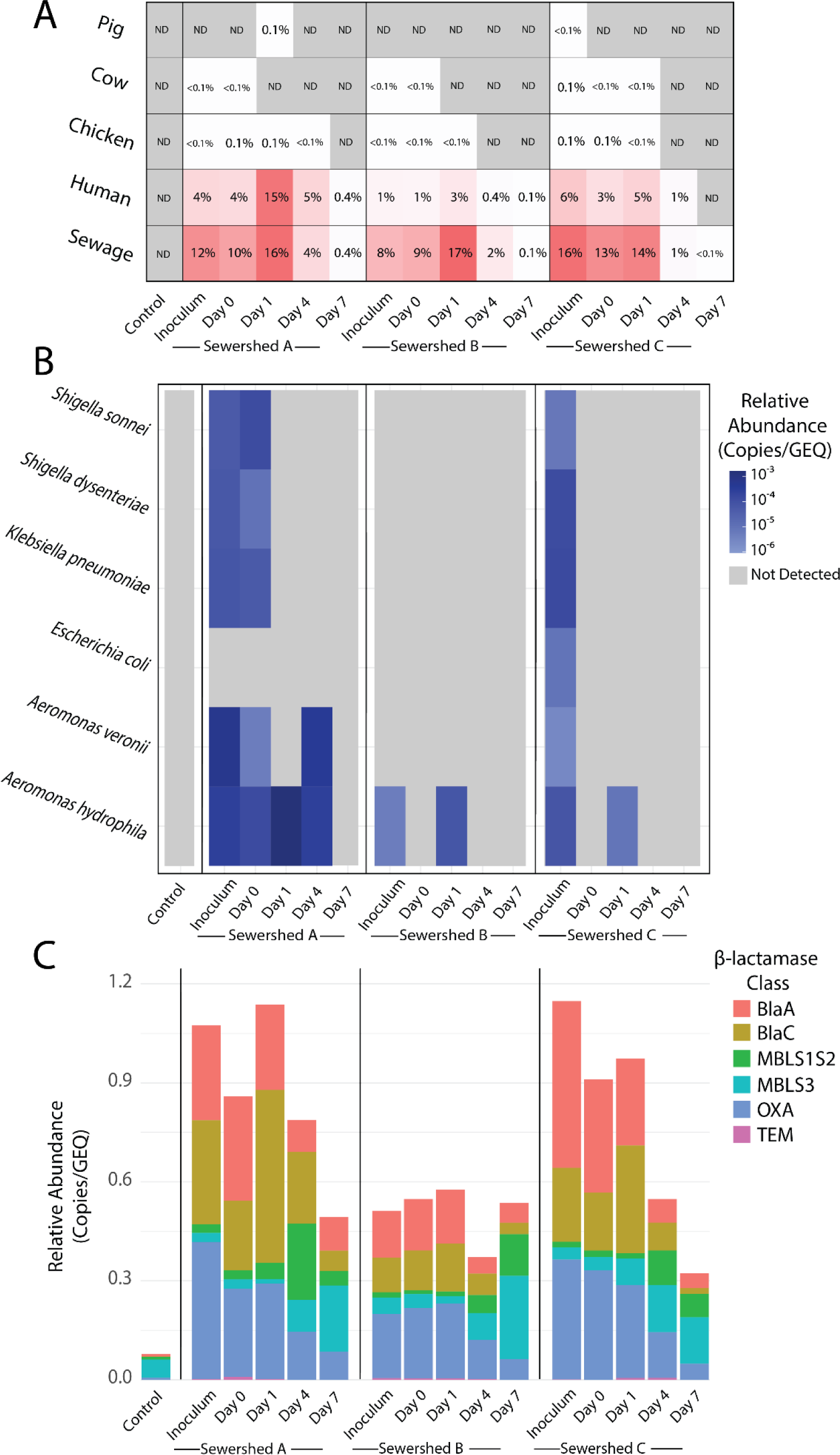
Abundance patterns of Source Tracking libraries, virulence factors and β-lactamase encoding genes across inoculum and timeseries metagenomes. All normalization was performed against genome equivalents (GEQ). **Panel A**: Source attribution and partitioning results based on reads mapped against MAGs curated for different fecal sources. Percentages represent estimates of the fraction of the prokaryotic population specifically belong to one of the fecal sources. **Panel B**: Virulence Factor (VF) gene abundance dynamics based on short reads mapping on experimentally verified VF reference nucleotide sequences (Figure key). **Panel B**: β-lactamase gene abundance dynamics across inoculum, timeseries and control metagenome based on Diamond --blastx searches of reads against reference ARG sequences and ROCker model filtering of the resulting matches. Relative abundance is calculated by normalizing the average sequencing depth of each gene to GEQ after ROCker filtering.

### Pathogen and Virulence Genes Assessment

To assess the ability of the metagenomic approach to provide insights into the health risk associated with bacterial pathogens introduced by sewage contamination during mesocosm operation, we recruited metagenomic short reads to 1076 pathogenic bacterial genomes recovered from the PATRIC webserver (**Supplement, Table S5**). Results revealed that 63, 38, and 129 pathogen genomes from sewersheds A, B, and C, respectively within the inoculum had sequencing depths at or above our established LOD after read mapping (see Methods, Eqn. 1) (**Supplement, Table S6**). In contrast, immediately following inoculation on day 0 many reference genomes were no longer detectable, with a total of 61, 25, and 20 pathogenic genomes detected from sewersheds A, B, and C, respectively. It should be mentioned, for many of these organisms, pathogenicity is a function of genotype (e.g., the *E. coli* pathotypes) and the methods used herein were developed for species-level detection and not optimized for distinguishing between closely related genotypes of the same species at low abundances (55).

Therefore, due to the low relative abundances of these pathogens that we observed and the need to assess the actual genetic content present within these populations, we examined the relative abundance of experimentally verified genes within the Bacterial Virulence Factor Database (VFDB) as proxies for key bacterial pathogens (**Figure 5, B**). The virulence signal within inoculum metagenomes primarily comprised those belonging to *Aeromonas*, *Klebsiella*, and *Shigella* pathogenic genera, consistent with the whole-genome detection results above. Sewage from both sewershed A and C appeared to have greater virulence factor signals compared to sewage from sewershed B, which had drastically lower detected levels of *Aeromonas* VFs and no detection of *Klebsiella*, *Shigella* or *Escherichia* VFs. Within the sewershed A and C timeseries, average virulence abundance was lower on day 0 than in the inoculum but quickly reached a maximum in 24 hours before substantially decreasing by day 4 and being below detection by day 7. The increase was primarily due to substantial increase in the abundance of *Aeromonas hydrophila* VFs. This trend was consistent among genes encoding for *hlyA* (hemolysin), *aerA* (aerolysin) and *act* (*Aeromonas* enterotoxin), essential cytotoxins for *Aeromonas* spp. pathogenicity, across the timeseries. Alignment of these three cytotoxin genes to the MAG representing Pop. 15 revealed that it likely carries a gene encoding for *hlyA* but *aerA* and *act* were either not binned with the draft genome or truly not carried by this population. Upon further inquiry, the closest matching entry on NCBI’s Genome database was *Aeromonas caviae* NZ_AP022214 (ANI = 98.0%), which represents a strain isolated from a Japanese wastewater treatment plant that has not been implicated in disease or designated as an obligate pathogen. Hence, to what extent the MAG identified represents a pathogenic or opportunities pathogenic population remains somewhat speculative.

### β-lactam Resistance Gene Assessment

Several classes representing the breadth of β-lactamase-encoding gene diversity were present in the metagenomes from all samples. The uninoculated lake water (control) sample showed very low abundance of β-lactamase encoding genes across each class (sum of classes was 0.078 total β-lactamase encoding genes/genome equivalent) – though a subset of metallo-β-lactamase encoding genes (MBLS3) were noticeably pronounced (0.06 gene copies/genome equivalent). In the inoculum samples, total observed β-lactamase signal was much greater in sewersheds A and C (1.07 and 1.14 total gene copies /genome equivalent, respectively) compared to sewershed B (0.51 total β-lactamase encoding genes/genome equivalent), but the relative contribution of each class was consistent, with genes encoding for BlaA, BlaC and OXA dominating. In contrast, by day 4 and to a greater extent by day 7, the frequency of genes encoding for BlaA, BlaC and OXA decreased consistently while those encoding for MBLs increased (**Figure 5**, C). Along with a shift in prominence of these β-lactamase gene classes, both sewersheds A and C showed steep decreases in the relative number of β-lactamase encoding genes/genome equivalent between day 0 and day 7. Sewershed C showed the same shifts in prominence between classes, yet total signal remained consistent with 0.55 and 0.54 total β-lactamase gene copies/genome equivalent on day 0 and 7, respectively.

## Discussion

### Sewershed Microbial Diversity

Collection systems represent a key component of modern sanitation infrastructure. Despite the importance of sewage as a reservoir for human pathogens and antimicrobial resistance genes, the sewage microbiome remains relatively understudied at the whole genome level. Our results indicated that the sewage samples we collected from three separate collection systems across the Atlanta Metropolitan region were dominated by what have been aptly named microbial “weeds” in literature and which we have observed as belonging to several sewage-associated populations which appear quite prolific (21, 22). Others have reported many of these populations are also present at high relative abundances within sewersheds spanning another urban landscape (72). Our resulting analysis expanded our understanding of these bacterial populations and their fate during a simulated contamination event by recovering representative draft genomes (MAGs) for these ubiquitous populations and tracking their abundances over time with controls in place for microbial load fluctuations. Specifically, we found primary sewage-associated populations to represent clades within the classes *Gammaproteobacteria* (e.g., *Acinetobacter*, *Aeromonas*), *Betaproteobacteria* (e.g., *Acidovorax*, *Rivicola*) and *Epsilonproteobacteria* (e.g., *Arcobacter*) (**Figure 1,A**).

These sewage-associated populations showed different preference for oxygen, appearing to span strict anaerobic, facultative, and aerobic metabolic phenotypes. Notably, the signal associated with these populations in the metagenomic datasets decayed non-uniformly during mesocosm operation, though the most persistent populations were aerotolerant, acetate-utilizing populations which contained genes related to aromatic degradation and/or nitrogen metabolism. Depending on additional inquiry, it may be possible to leverage the ratio between abundances of anaerobic and aerobic (or facultatively anaerobic) sewage-associated populations in future work for inferring the date of pollution events linked to sewage contamination. For all 15 populations described here, their linear relationship with HF183 and Enterococci had a combined R^2^ of 0.6 (**Figure 2**), revealing overall consistent results for different markers under the conditions tested here. However, these correlations were drawn from the limited number of mesocosm incubations and *in situ* population dynamics are likely to differ according to varying environmental and biological factors which were not controlled for herein.

Our dataset is of limited size and scope considering that, on a global scale, we examined sewage from collection systems in essentially equivalent geographies. The assortment of sewage-associated populations described here, although ubiquitous across the sewersheds we sampled, likely maintain differing prevalence across time or space. Furthermore, many draft genomes we produced are not complete,so further work will be needed to establish a more practical view of both the range of these populations and their genomic content and diversity. Yet, we see advancing our knowledge of sewage-associated populations as a potential contribution towards newly developing forensic approaches that help monitor, manage, and repair these essential infrastructures (73). For example, we observed several highly abundant populations with a range restricted to only one or two of the three sewersheds. It would be important to gauge whether populations (or genotypes within a population) exist that are specific to individual sewersheds, and how the physicochemical characteristics (e.g., municipal vs. industrial waste, flow rates) of different waste streams might drive the formation of these distinctions. Further inquiry in this direction may also lead to strategies for resolving source attribution inquiries when multiple collection systems with differing catchment compositions are all possible sources of contamination in the same water environment.

### Source Attribution and Partitioning with Host-Specific Genomic Libraries

Several of the abundant sewage-associated populations identified above appear closely related (likely at the species level) to members of human or chicken fecal microbiomes (**Figure 4**). Regardless, it appears populations specific to municipal sewage likely exist and represent a contingent of the sewage microbiome which – if better catalogued – may be useful for identifying and quantifying sewage pollution in natural ecosystems independent of human-associated markers. We have demonstrated, through a proof-of-concept workflow, the capacity for read mapping metagenomic datasets to host-specific genomic libraries for performing both source attribution and partitioning (**Figure 5, A**). Although source attribution has been well-developed in existing metagenomic approaches, no metagenomic methods have been developed which are also capable of simultaneous source partitioning. Here, we have performed source partitioning for each entry within a host-specific genomic library to a metagenome’s GEQ, which yields a relatively easy-to-interpret metric describing the percentage of genomes within a metagenome that belong to a given host or source specific library. We see this as a promising avenue for metagenomic-based MST methods and believe the approach could eventually be utilized in the field pending further testing and refinement by mixed input experiments.

### β-lactamase Encoding Genes Surveillance

Additionally, we leveraged our metagenomes to survey for β-lactamase genes across the inoculum and timeseries. The abundance of β-lactamases across the inoculum samples was substantially higher (7-15 times) compared to the control (**Figure 5**). This result was consistent with both our expectations and the literature regarding heightened ARG abundance within collection systems (74). Specifically, others have reported substantial abundances of β - lactamase OXA genes on both *Campylobacteraceae* and *Aeromonadaceae* clades in sewage (75). Indeed, the abundance of reads belonging to β-lactamase encoding genes, especially of the OXA-encoding class, were the most abundant in the inoculum and early time points where these sewage-associated clades (e.g., Pop.01, Pop.19) persisted in the lake water. Overall, these results indicated that sewage contamination imparted a substantial and lasting increase to the abundance of genes encoding β-lactamases even after 7 days following the contamination event. More work is needed to elucidate the genomic context of this increased β-lactamase encoding gene abundance (e.g., whether they belong to or have been transferred to organisms capable of driving clinically relevant cases of antimicrobial resistance). Nonetheless, our results allow for a quantitative view of the abundance of these genes relative to the natural environment where the freshwater used in the mesocosm incubations originated, which is relevant for assessing the associated public health risk.

### Pathogen and Virulence Gene Surveillance

Importantly, although Sewershed A and B showed what appears to be similar concentrations of human input according to HF183 concentrations within the inoculum (**Supplement Figure S1**), the pathogen detection results revealed via the sequence data were quite varied (**Figure 4B**, **Supplement Table S6**). Results from both read mapping to bacterial pathogen genomes and the experimentally verified VFDB collection were consistent in suggesting that bacterial virulence was more elevated in the Sewershed A inoculum compared to Sewershed B. This contrast between sewersheds with equal human marker concentrations yet apparently unequal bacterial pathogen load illustrates how shotgun sequence data can facilitate perspectives on the actual co-variance of marker and pathogen. Yet these insights clearly depend on sufficient sequencing effort and/or relatively high pathogen concentrations to avoid the possibility of false negative results.

In particular, the estimated smallest detectable population size associated with our analysis and sequencing effort ranged between approximately 2E+05 to 1E+02 cells/mL based on qPCR-based cell count normalization and the sequencing effort applied (Methods, **Supplement Table S2**). Approaches for estimating analytical LOD within metagenomic based analysis remain rare within the literature especially as it relates to work done in the environment as opposed to clinical settings (76, 77). Yet, the concept of detection and quantification limits in metagenomics is a major challenge to its thorough incorporation into environmental monitoring approaches because 1) it is necessary to track biomarkers or pathogens down to quite low relative abundance in the field (i.e., <1E-09 target basepairs/total basepairs), and 2) leveraging extraordinary sequencing effort is currently expensive and not practical when limitations of expertise and computational resources exist. Our approach provides the means to establish theoretical analytical LOD for metagenomic analyses based on sequencing effort which is useful for determining and interpreting the meaning of “non-detects”.

Using AGS and total cell density estimates within the inoculum, we estimate approximately 3.5Tb of sequencing effort is necessary for detecting a population with concentration of 1E+02 cells/mL within the high microbial loading conditions observed in the inoculum. In contrast, following the decline in cell density and increase in AGS across the timeseries, the estimated sequencing effort required to detect a population of 1E+02 cells/mL drops to 10Gb in day 7 conditions. Therefore, our approach and results reported here for sequencing effort estimation may be helpful for informing the planning and execution of future environmental monitoring work utilizing metagenomic approaches (**Supplement Table S7**). Though, crucial to note is the fact that our approaches for analytical LOD, and sequencing effort estimation assumes unbiased sequencing and does not consider sampling or processing recoveries – where the latter limitation is obviously broadly applicable to all molecular methods. Total detection limits, in the context of analytical limits as well as both sequencing bias and sampling/processing recoveries, will be important caveats to consider for future metagenomic workflows aiming to surveil pathogens in sewage collection systems or their releases into the environment (78).

While we do not envision that PCR and culturing will be replaced by metagenomics for routine monitoring because the former techniques are cheaper, easier to analyze, and have a greater dynamic range of detection – our results do show how metagenomics can provide unique insights into sewage pollution events such as differential pathogen content of different sewage or possibly distinguishing between different sewersheds potentially contributing to contamination. Further, we have shown how metagenomics could track a broad range of population sizes – about six orders of magnitude (from about 1E+01 to 1E+07 cells/mL) – which is adequate for certain applications and/or high-volume pollution events.

### Conclusions

Our efforts have shown how metagenomic datasets can provide insights on multiple questions critical to environmental monitoring and water quality: pathogen detection, source attribution and partitioning, and ARG persistence in the environment. In our view, confident and direct detection of pathogens within metagenomic datasets will remain primarily a logistical challenge due to the large amount of sequencing effort required to reliably detect bacterial pathogens at concentrations that are very low yet still quite relevant to protecting public health. Thus, when performed alone, metagenomic approaches are unlikely to be the most prudent technology for routine monitoring and directly informing health risks associated with sewage contamination, especially when pathogen or virulence genes are at these relatively low abundances (e.g., below 1E+02 features/mL). This issue is also compounded by the large contribution of non-bacterial pathogens (e.g., viruses and protozoa) to illness risk in contaminated waters. In contrast, metagenomic approaches are increasingly poised to resolve questions related to source attribution and partitioning by improving our understanding (and the size of our databases) of the genomes maintained by source-specific microbial populations.

## Acknowledgments

The authors would like to thank the Cobb County Water System, Gwinnett County Department of Water Resources, and the City of Atlanta Department of Watershed Management for assistance with this work. This research was supported in part through research cyberinfrastructure resources and services provided by the Partnership for an Advanced Computing Environment (PACE) at the Georgia Institute of Technology, Atlanta, Georgia, USA. This work was supported by the US National Science Foundation, award numbers 1511825 (to J.B and K.T.K) and 1831582 (K.T.K.) and the US National Science Foundation Graduate Research Fellowship under grant number DGE-1650044 (to B.S.). The funding agencies had no role in the study design, data collection and analysis, decision to publish, or preparation of the manuscript.

