## Supplement Figures for "Novel municipal sewage-associated bacterial genomes and their potential in source tracking"

for article entitled

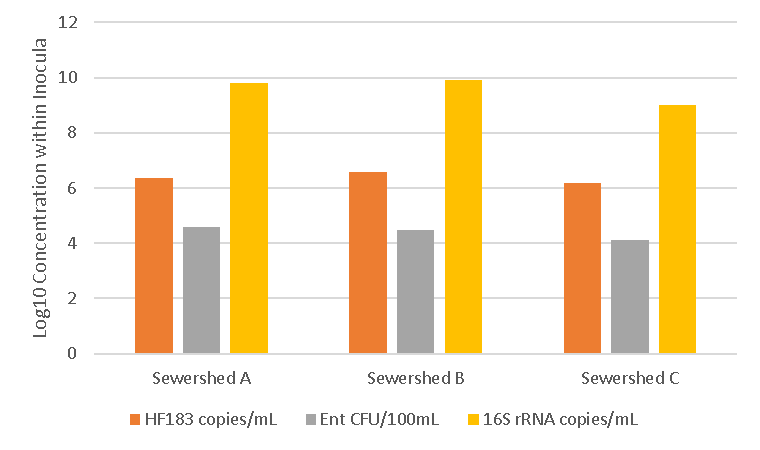

**Figure S1. Average qPCR (HF183 and GenBac 16S rRNA assays) and culture-based (EPA-1600) counts between sewage inocula samples, revealing unequal microbial load and human fecal input across sewersheds.**

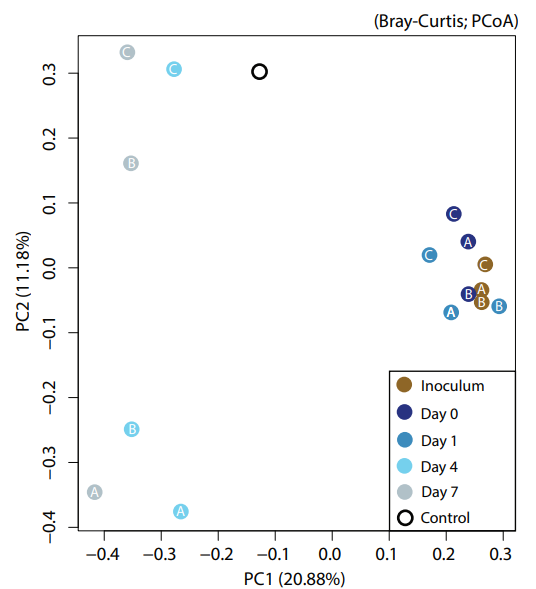

**Figure S2: Bray-Curtis distance matrix calculated and visualized using simka software on the metagenomic short reads of the control, inocula and mesocosm samples.** Note that data points early in the operation of the mesocosms (day 0, day 1) remained very similar to the inocula compared to the control or later mesocosm time points.

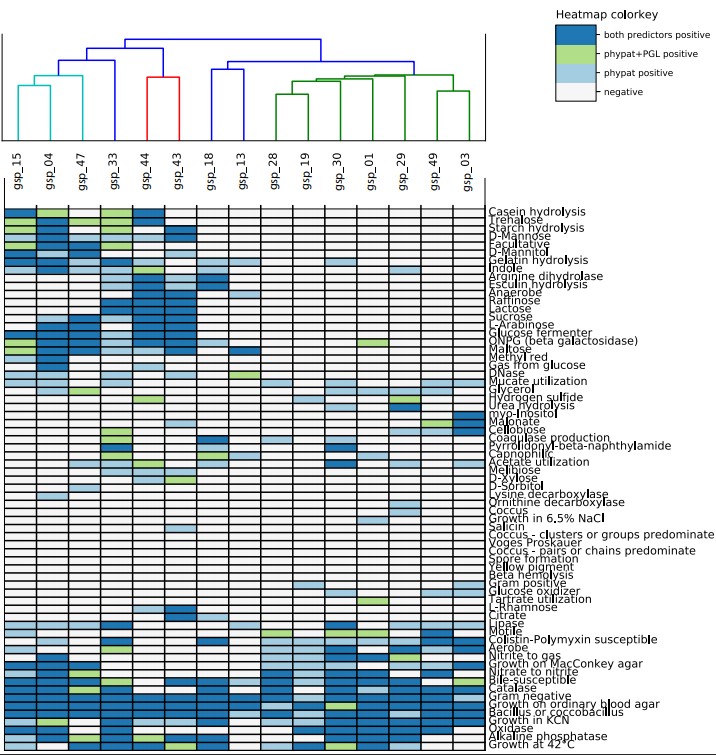

**Figure S4: Traitar phenotyping results with clustering by MAG functional pathway relatedness (top) and heatmap indicating annotation consensus (on the right).**

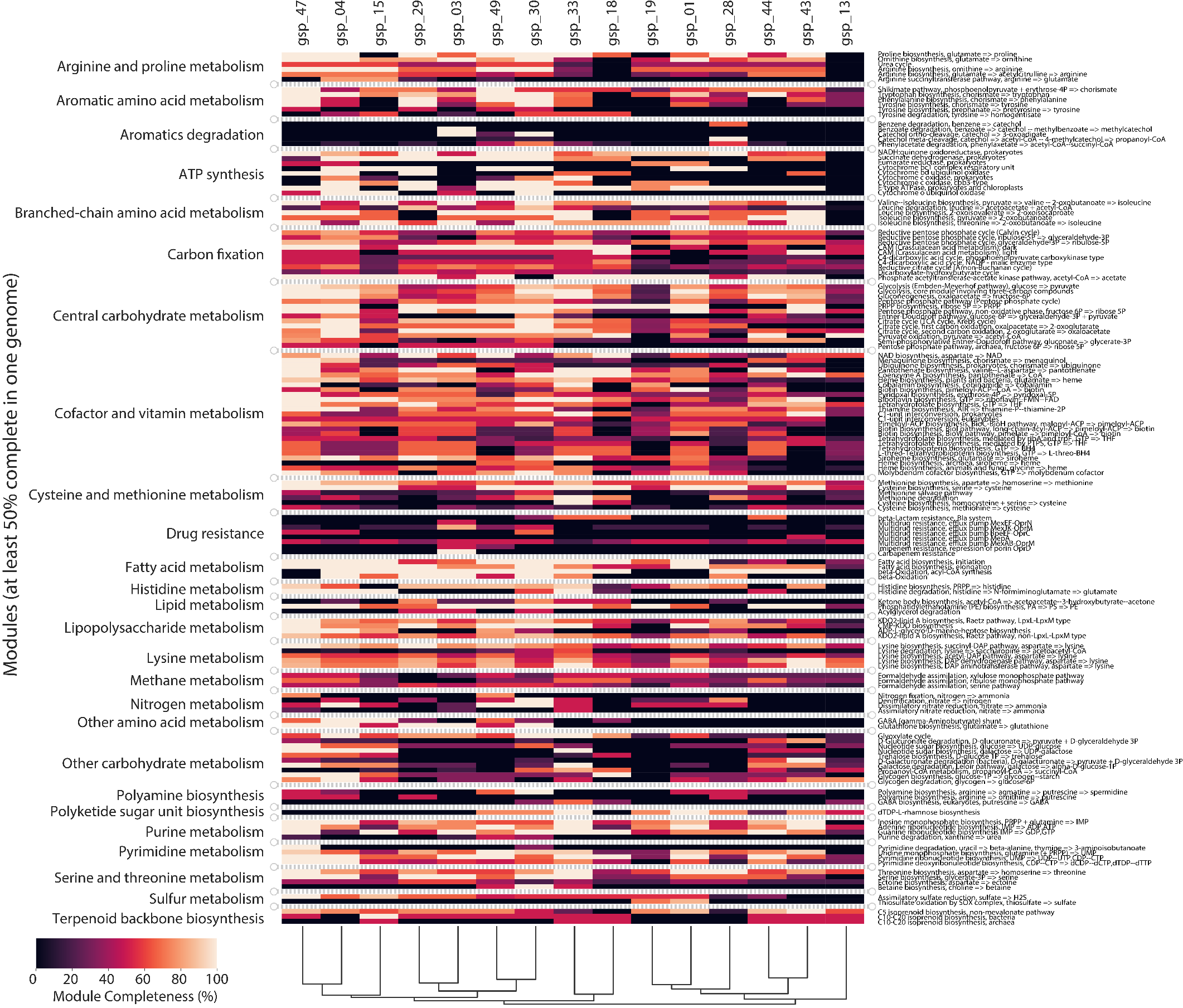

**Figure S5: Heatmap showing the relatedness of MAG based on their predicted functional gene pathways.** Pathways were predicted using the MicrobeAnnotator and results are sorted by both MAG relatedness (columns) and module similarity (rows). Module completeness was predicted by MicrobeAnnotator based on the software’s ability to identify sufficient genes to complete a functional pathway.
